# Embryo-to-embryo variability in RNAi knockdown efficiency of *dKDM5/lid in Drosophila melanogaster*

**DOI:** 10.1101/2021.04.29.442033

**Authors:** Ashley Albright, Michael Eisen

## Abstract

We used the maternal-Gal4 shRNA system to knock down expression of *dKDM5/lid* in *Drosophila melanogaster* embryos, and analyzed the efficacy of the knockdown by qRT-PCR. Although average relative expression of *lid* was significantly lower in knockdown conditions compared to the driver-only control, we observed a wide and overlapping range of relative gene expression between individual control and knockdown embryos.

## Introduction

The maternal-Gal4 shRNA system is widely used to knock down gene expression in Drosophila embryos^1,2^. We carried out several experiments using this system to knockdown *dKDM5/lid* (subsequently referred to as *lid*) with the goal of comparing gene expression of *lid* knockdown embryos to a driver-only control via single-embryo RNA-sequencing. Before sequencing, we used qRT-PCR to characterize the efficacy of knockdown and found a surprising overlap of *lid* expression between individual control and knockdown embryos.

## Methods

### Fly Husbandry

All stocks and crosses were fed a molasses-based diet prepared by the UC Berkeley Media Prep Facility and maintained at 25 °C. All UAS-shRNA lines were obtained from the Bloomington Drosophila Stock Center. RNAi was done by crossing nanos-Gal4; HisRFP virgin females to UAS-shRNA males from the Transgenic RNAi Project as described previously^2^. Flies were placed into collection cages 3-4 days after hatching. Cages were fed with yeast paste made of Red Star yeast pellets and water on grape juice agar plates to feed the caged flies every day.

### Single-embryo RNA extraction and qRT-PCR

Cages were cleared for 30 minutes to 1 hour prior to a collection for 1.5 hours and aged for 70 minutes prior to extraction. Embryos were dechorionated by agitating in 50% bleach for 3 minutes. Up to six stage 3 embryos at a time^3^, limited by the number of dounces available, were sorted by visualizing them under a dissection microscope and transferring the embryos with a paintbrush into separate dounces containing 1 mL Trizol (Invitrogen). RNA from single embryos was extracted as described previously^4^; however, we modified homogenization by using a dounce instead of a needle. Embryos were homogenized in Trizol by douncing 20 times with a loose pestle (A) and 10 times with a tight pestle (B). At this point, embryos were either frozen in Trizol at −80°C or we continued with the manufacturer’s instructions. RNA was further cleaned using Turbo DNase (ThermoFisher) according to the manufacturer’s instructions to remove any contaminating DNA, and quantified using a Qubit 2.0. Quality of RNA was assessed on a Bioanalyzer using the RNA 6000 Pico Kit (Agilent). Acceptable samples were determined by eye, as *Drosophila* ribosomal RNAs run too close to each other for an accurate RIN score.

Level of *lid* RNA in 8 control, and 8 knockdown embryos of each separate shRNA line was measured using the Invitrogen SuperScript™ III Platinum™ SYBR™ Green One-Step qRT-PCR Kit using the primers in Table 1. Primers were designed using FlyPrimerBank^5^. In an attempt to alleviate technical variability in cDNA synthesis or measurements by the machine across plates, we conducted qRT-PCR in 8 embryos per condition with 2 technical replicates each such that everything was run on one plate.

**Table 1.**
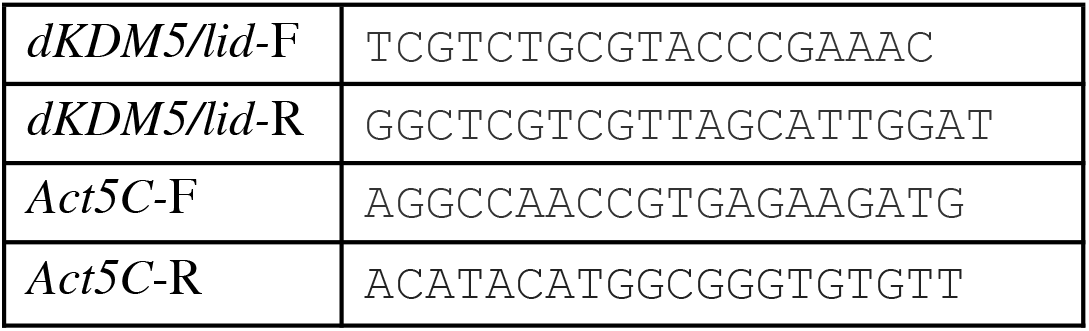
Primer sequences for qRT-PCR.

Samples were run on a Roche Lightcycler 480 at 50°C for 3 minutes, 95°C for 5 minutes, followed by 40 cycles of 95°C for 15 seconds, 55°C for 30 seconds, and 72°C for 60 seconds. The reaction was held at 40°C for 1 minute followed by melting curve analysis to ensure amplification was of a single product.

Using R, we calculated relative *lid* expression using the 2^-ΔΔCp method^6^ with *Act5C* as a control. Figures were generated using ggplot2^7^, and full code is available at: https://github.com/aralbright/2021_AAME_qPCR.

## Results

We used FlyPrimerBank to design our primers and the authors validated many qPCR primers and RNAi lines in embryos using bulk RNA extractions^5^, so we decided to first determine the efficacy of RNAi knockdown by mimicking the same with our single embryo extractions. To mimic collecting RNA in bulk, we calculated the average Cp value of the individual samples prior to calculating 2^-ΔΔCp (subsequently referred to as pseudo-bulking). Knockdown appears efficient relative to the driver only control with relative expression at 0.11 and 0.22, for 35706 and 36652 respectively (Figure 1A). To obtain relative gene expression values for individuals, we calculated 2^-ΔΔCp for each embryo relative to the average control embryo. In this case, average relative expression of control, 35706, and 36652 samples are 1.05, 0.38, and 0.55 respectively (Figure 1B). While these averages are higher than those shown in Figure 1A, in both cases *lid* expression is lower in the population of knockdown embryos than in the driver-only control.

**Figure 1.**
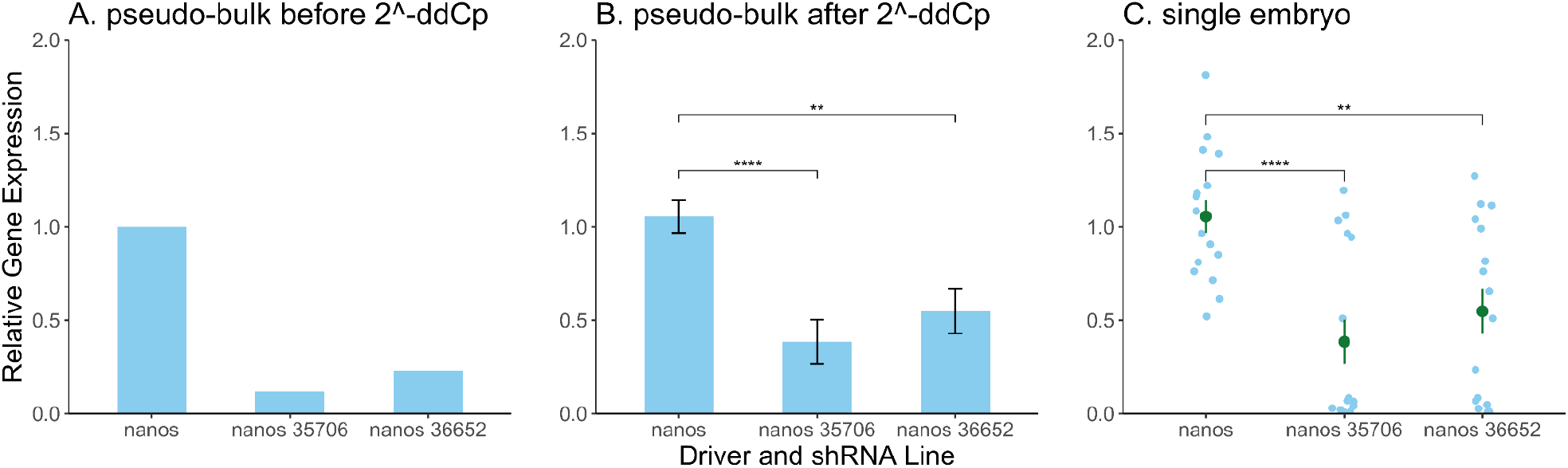
Relative gene expression of *lid* as measured by qRT-PCR in Stage 3 embryos plotted by **(A)** averaging each group prior to calculating 2^-ΔΔCp, **(B)** after calculating 2^-ΔΔCp, and **(C)** considering each embryo as an individual sample. The difference between control and knockdown embryos as shown in Figure 1B and Figure 1C is statistically significant with adjusted p-values of 0.00029 and 0.0038, for lines 35706 and 36652 respectively. Error bars indicate standard error.

By visualizing the distribution of individual values instead of their mean (Figure 1C), we revealed an overlap between individual control and knockdown embryos. The overall average and statistical significance values in Figure 1C are the same as Figure 1B; however, the range of overlap between the control and both RNAi conditions as depicted in Figure 1C indicates possible biological and/or technical variance of expression level within each condition that is not clear from Figure 1B.

To further describe the variability shown in Figure 1C, we show the data colored by each individual embryo within each condition in Figure 2. While a difference of relative gene expression between some technical replicates indicates the presence of technical variability, 3/8 embryos using line 35706 have greater than zero expression and 5/8 embryos for 36652.

**Figure 2.**
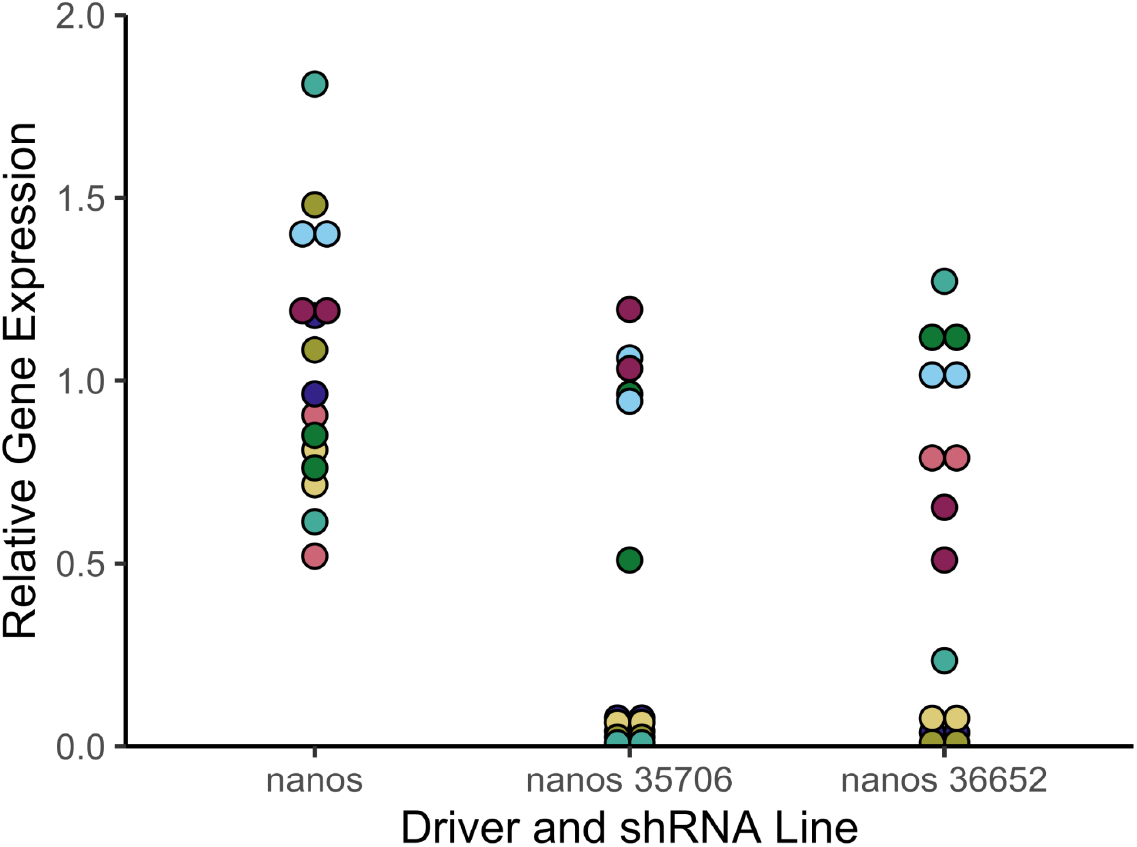
Relative gene expression of *lid*. Colors representing individual embryos within each condition.

We have demonstrated that some embryos retain expression upon knockdown of *lid*, and relative expression values overlap widely with the driver-only control. This, or masking the variability by collecting samples in bulk, might complicate downstream experiments.

## Acknowledgements

We thank members of the Eisen Lab for feedback. AA was supported by an NIH Training Grant (T21 GM 007127) and the National Science Foundation Graduate Research Fellowship Program. The work was also supported by a Howard Hughes Medical Institute Investigator award to ME.

